# The RNF220 domain nuclear factor Teyrha-Meyrha (Tey) regulates the migration and differentiation of specific visceral and somatic muscles in *Drosophila*

**DOI:** 10.1101/2022.11.18.517102

**Authors:** Manfred Frasch, Afshan Ismat, Ingolf Reim, Jasmin Raufer

## Abstract

The development of the visceral musculature of the *Drosophila* midgut encompasses a closely coordinated sequence of migration events of cells from the trunk and caudal visceral mesoderm, respectively, that underlies the formation of the stereotypic orthogonal pattern of circular and longitudinal midgut muscles. Our current study focuses on the last step of migration and morphogenesis of the longitudinal visceral muscle precursors derived from the caudal mesoderm. We show that these multinucleated muscle precursors utilize dynamic filopodial extensions to migrate in dorsal and ventral directions over the forming midgut tube. The establishment of maximal dorsoventral distances from one another and subsequent alignment with their anteroposterior neighbors leads to the equidistant coverage of the midgut with longitudinal muscle fibers. We identify Teyrha-Meyhra (Tey), a tissue-specific nuclear factor related to the RNF220 domain protein family, as a crucial regulator of this process of muscle migration and morphogenesis that is further required for proper differentiation of the longitudinal visceral muscles. In addition, Tey is expressed in a single type of somatic muscle founder cell in each hemisegment. Tey regulates the migration of this founder cell and is required for the proper pathfinding of its developing myotube to specific myotendinous attachment sites.

## Introduction

In metazoans, proper organogenesis relies on precisely controlled cell migration events and dynamic cell shape changes that sculpt each organ. In most cases, several different cell types participate in these events and must closely coordinate their dynamic behaviors in order to build a proper organ. Although past and ongoing research has uncovered a large body of information on the regulation of cell migration and cellular morphogenesis during organ formation (Aman and Piotrowski, 2010; Scarpa and Mayor, 2016; Stock and Pauli, 2021), we currently are still lacking a full understanding of how most organs are being formed during development.

In *Drosophila*, one organ that has been used as a favorable model for dissecting these events is the midgut, and in particular, the musculature that ensheathes it. In its fully differentiated state, at the end of embryogenesis as well as in the adult fly, the musculature around the tube of endodermal cells of the midgut consists of a highly ordered arrangement of two different types of muscle fibers. One type, the circular visceral muscles, consist of elongated, binucleated syncytial cells that are oriented exactly in dorsoventral directions. In this manner, they form semi-circles around either side of the midgut that are attached to their contra-lateral equivalents along the dorsal and ventral midlines, thus collectively encircling the endodermal tube. The second type, the longitudinal visceral muscles, consist of multinucleated muscle fibers that extend in anteroposterior directions along the midgut tube and, thus, are oriented precisely orthogonally to the circular visceral muscles. The longitudinal muscles largely sit on top of the circular ones, although they are partially interwoven with them (Campos-Ortega and Hartenstein, 1997; Lee et al., 2005; Schröter et al., 2006). Unlike the circular muscles, the longitudinal muscles maintain significant distances from their neighbors and thus form an evenly-spaced arrangement of parallel muscle fibers along the midgut. It is a very intriguing question how this precisely orthogonal network of muscle fibers is being assembled during development, especially in light of the fact that the gut tube does not yet exist when the visceral muscle progenitors are born and that the progenitors of the two different visceral muscle types have completely different origins in the early mesoderm.

Both the founder and fusion-competent cells of the circular visceral muscles (CiVMu’s) are derived from the trunk visceral mesoderm (TVM), which is being specified as 11 segmental cell clusters by ectodermal patterning signals along the dorsal mesoderm of the trunk region (Azpiazu and Frasch, 1993; Lee et al., 2005). At the beginning of germ band retraction, these TVM clusters rearrange to form a continuous narrow band of TVM cells along the a/p axis on either side of the trunk underneath the somatic mesoderm. This structure is then used as a track for the migration of cells of the anterior and posterior endoderm, which ultimately meet in the middle and subsequently extend in dorsoventral directions to ultimately form the endodermal portion of the midgut tube (Azpiazu and Frasch, 1993; Reuter et al., 1993; Tepass and Hartenstein, 1994). During the same period, each CiVMu founder cell of the TVM undergoes myoblast fusion with a single fusion-competent cell from the TVM to form binucleated circular visceral muscle precursors (CiVMp’s), which then elongate dorsoventrally and line up next to each other into a palisade-like arrangement (San Martin et al., 2001). Continued dorsoventral elongation of the CiVMp’s in concert with the endoderm leads to the circular visceral musculature of the midgut.

In contrast to the TVM that is generated all along the a/p axis of the embryonic trunk, the progenitors of the longitudinal midgut muscles are derived from a small region at the posterior end of the early mesoderm, termed caudal visceral mesoderm (CVM) (Georgias et al., 1997; Nguyen and Xu, 1998). The CVM is specified by the bHLH transcription factor HLH54F that becomes expressed in this region during the late blastoderm stage (Ismat et al., 2010). Cells from this cluster undergo several consecutive migratory events towards the anterior, thereby spreading out along the prospective midgut region and forming the founder cells of the longitudinal midgut muscles (Kusch and Reuter, 1999; Lee et al., 2005). First, after splitting into two bilateral clusters, each of these migrates as a collective towards the respective posterior-most cluster of the TVM on either side. Second, cells from these bilateral clusters of CVM begin streaming anteriorly on either side of the embryo as a more loosely organized collective (Sun et al., 2020). This migration occurs along precise tracks both at the dorsal and the ventral margins of the bands of TVM and in close contact with them. During this migration, the CVM cells become the founder cells of the longitudinal visceral muscles, and upon their fusion with the remaining fusion-competent myoblasts from the TVM and after spreading evenly along the length of the TVM bands they form the longitudinal muscle precursors (LVMp’s) (San Martin et al., 2001).

The anterior long-distance migration of the CVM cells recently has received considerable attention and some of the mechanisms guiding it have been uncovered (reviewed in Sun et al., 2020). The requirement for the TVM as a migration substrate upon stage 12 was demonstrated by genetically ablating the TVM (Zaffran et al., 2001; Reim et al., 2012). Several extracellular matrix (ECM) components likely laid down by the TVM, as well as specific signaling molecules that are released by it, have been found to promote CVM migration along the TVM. In particular, the expression of αPS2 integrin (encoded by *if*) in the TVM is instrumental in the normal deposition of Nidogen in the ECM along the TVM, and mutations affecting either of these two components display significant delays in CVM migration (Urbano et al., 2011). Mutations in lamininW subunits cause even more severe CVM migration defects, which implicated lamininW as one of the key components in the ECM along the TVM to promote anterior CVM migration. αPS1 integrin (encoded by *mew*) being expressed in the migrating CVM cells makes a similar contribution to the migratory efficiency as αPS2 in the TVM (Urbano et al., 2011). Other components interacting with the ECM or being part of it, including the secreted metalloproteinase AdamTS-A, the chondroitin sulfate proteoglycane Kon-tiki and the heparan sulfate proteoglycane Trol, were also found to promote anterior CVM migration (Ismat et al., 2013; Trisnadi and Stathopoulos, 2014; Hamilton et al., 2022). In addition to facilitating direct interactions between the TVM substrate and the migrating CVM cells, it is conceivable that some of these ECM components also play roles in regulating the distribution of FGF ligands, which have been identified as essential signaling molecules in the regulation of anterior CVM migration. These ligands, Pyramus (Pyr) and Thisbe (Ths), are expressed in the TVM (and, in the case of Pyr, additionally in the adjacent endoderm), whereas their dedicated FGF receptor Heartless (Htl) is highly expressed in the migrating CVM cells (Mandal et al., 2004; Reim et al., 2012). Importantly, a series of genetic experiments has demonstrated that FGF signaling is required for proper pathfinding of the migrating CVM cells, for them to remain attached to the TVM during their migration, and for securing the survival of the migrating CVM cells (Kadam et al., 2012; Reim et al., 2012; Macabenta et al., 2022).

Very interesting migration events take place also in the somatic mesoderm and are essential for the formation of a properly patterned skeletal muscles. These migrations involve the directed migration of somatic muscle founder cells relative to one another and the epidermis (e.g., Dohrmann et al., 1990), which upon myoblast fusion is followed by distinct pathfinding processes of the leading edges of the muscle precursors towards their respective myotendinous junctions (Schnorrer and Dickson, 2004). Whereas the mechanisms underlying the founder cell migrations are completely unknown, several of those involved in myotube pathfinding have been determined. Some may be similar to the ones described for CVM migration, which may be the case for the roles of Laminins and Kon-tiki (Estrada et al., 2007; Schnorrer et al., 2007; Wolfstetter and Holz, 2012; Perez-Moreno et al., 2017; Perez-Moreno et al., 2021). Others appear specific to myotube migration, particularly the signaling components guiding the myotubes to their muscle attachment sites. An important signal known to be released from the tendon progenitors and to attract the myotubes towards them is Slit, which activates Robo receptors on the surface of the extending myotubes (Schweitzer et al., 2010).

In contrast to the anterior migration of the CVM cells and their derivative longitudinal muscle founder cells, the subsequent events of the dorsal and ventral migration of the longitudinal muscle precursors and their coordinated alignment along the length of the midgut have not been investigated extensively. Herein, we show that these muscle precursors perform these migrations as syncytia ultimately containing ~ 6 nuclei, which are decorated on their surface by long and highly dynamic filopodia that extend towards their neighbors and their migration substrate. These mutual interactions, as well as the formation of anteroposterior polarities of these muscle precursors and the establishment of anterior/posterior contacts among them, are likely to be instrumental in the formation of the regular pattern of evenly-spaced longitudinal muscle fibers covering the midgut. Furthermore, we identify the nuclear protein Teyrha Meyrha (Tey), a diverged member of the RNF220 family of ubiquitin ligases, as an important factor in the coordinated dorsoventral migration and proper morphogenesis of longitudinal visceral muscle precursors both during embryogenesis and during metamorphosis. In addition to the specific expression and function of Tey in the migrating longitudinal visceral muscle precursors and their progenitors, we show that Tey is also expressed in the founder cells and precursors of a single type of somatic muscle, termed M12 (aka, VLM1), and is important for the correct migration of these founder cells as well as for the proper pathfinding of the M12 muscle precursors to target their native tendon cells at the anterior and posterior segment borders.

## Materials & Methods

### *Drosophila* stocks

Offspring from *Drosophila* crosses were grown at 25 °C except for *UAS/Gal4* crosses, which were developed at 29 °C. The following stocks were either described in our previous publications or were obtained from the Bloomington *Drosophila* Stock Center (unless denoted otherwise): *Fas3-nGFP* (Jin et al., 2013); *HLH54Fb-GFP* (Ismat et al., 2010); *HLH54Fb-cytoRFP* (aka, P{HLH54F.LVM-RFP}16c; Hollfelder et al., 2014); *UAS-His-RFP* (Emery et al., 2005); *tey^5053A^* (aka, *tey-Gal4*; Lopez, J., 1998.11.24, pers. comm. to FlyBase; Inaki et al., 2010); *UAS-tey* (Inaki et al., 2010); *UAS-lacZ* (aka, *P{UAS-lacZ.B}melt^Bg4-2-4b^*; Brand and Perrimon, 1993); *UAS-Lifeact-GFP* (Hatan et al., 2011) (provided by F. Schnorrer); *bap3-RFP* (aka, *P{bap-RFP.3}7*; Reim et al., 2012)*; TM6 Dfd-EYFP* (Le et al., 2006); *caps-lacZ* (Shishido et al., 1998) (provided by A. Nose); *Df(3L)A23* (Cooper et al., 2010) (provided by J. Kennison); *Df(3L)ED228* (Ryder et al., 2004); *RRHS59-lacZ* (Knirr et al., 1999); *twi-Gal4* (aka, *P{GAL4-twi.2xPE}2* (provided by G. Schubiger); *HN39org-1-GAL4 & S18org-1-RFP* (Schaub et al., 2012); *UAS-apoliner9* (aka, P{w[+mC]=UAS-Apoliner}9 (Bardet et al., 2008); *nos-Gal4VP16 UAS-cas9* (BL-54593); *alphatub-piggyBacK10}M6; MKRS/TM6B,Tb* (BL-32070).

### Generation of CRISPR/Cas9 mutants for *tey*

*tey^Δ.M2-1^* and *tey^Δ.M1-11^* were generated by utilizing the system described in Port and Bullock (2016) and www.crisprflydesign.org to create UAS inducible t::gRNA array expressing transgenes. The amplifications of three PCR fragments containing four gRNA sequences (Fig. S5) with the respective primers and pCFD6 as a template (Addgene plasmid # 73915; from Simon Bullock), followed by Gibson assembly with the NEBuilder® HiFi DNA Assembly Cloning Kit (New England Biolabs), was performed as described in Schaub et al. (2019). The obtained *tey*-pCFD6-4 plasmid containing the four *tey* guide RNAs under the control of UAS sequences were injected for insertion into *AttB40* (BestGene Inc, Chino Hills, CA). Homozygous transgenic lines were crossed with *nos-Gal4VP14 UAS-cas9* and balanced flies from established lines were tested by PCR and sequencing for CRISPR/Cas9-induced deletions in *tey*.

*tey^ΔRNF.sfGFP^* was generated by replacing genomic *tey* sequences between the beginning of exon 4 and the end of the RNF domain-encoding sequence with the sfGFP-3xP3-TTAA-DsRed cassette from pHD-sfGFP-ScarlessDsRed (DGRC Stock 1365; https://dgrc.bio.indiana.edu//stock/1365; RRID:DGRC_1365; donor: Kate O’Connor-Giles) (with sfGFP being in frame) through homology-directed repair, followed by the scarless removal of the DsRed marker cassette flanked by piggyBac transposon ends via piggyBac transposase (Fig. 3).

For generating the donor plasmid, a 1.04 kb genomic 5’ homology arm, a 1.3 genomic 3’ homology arm, and a 2.4 kb fragment from pHD-sfGFP-ScarlessDsRed containing the sfGFP-3xP3-TTAA-DsRed cassette were PCR-amplified with the primers shown in Fig. S5. The three fragments were joined with one another and with the 4 kb SgrAI/PstI-cut vector backbone from pHD-sfGFP-ScarlessDsRed by Gibson assembly (see above) to obtain the *tey*-sfGFP-3xP3-TTAA-DsRed donor plasmid. For generating the guide RNA plasmid, PCR fragments with the respective primers containing the gRNA sequences (see Fig. S5) followed by Gibson assembly was done as described above, but in this case using pCFD5 with the corresponding protocol (www.crisprflydesign.org; Addgene plasmid # 73914; from Simon Bullock). The donor vector and gRNA plasmid (pCFD5-4) were co-injected into *vas-Cas9* embryos (BL-55821; BestGene Inc, Chino Hills, CA) and the single RFP^+^ line obtained was balanced. The DsRed-containing sequences between the TTAA-containing piggyBac recombination sites were then removed via crosses with a *alphatub-piggyBac*-containing line. Sequencing of the mutated locus confirmed the intended fusion of the Tey N-terminal peptide with sfGFP in conjunction with the deletion of sequences encoding in the central portion of Tey, including the RNF domain, starting from 3L:19653900 (Fig. S1, Fig. S5), as well as the scarless removal of DsRed. As designed, the sfGFP ORF continued in frame with ^817^V in the Tey C-terminus However, instead of continuing to the stop codon as intended, an apparent recombination error caused the sequence to continue after the second of the last codon (3L:19650976, ^821^M) with *tey* sequences starting from 3L:19651689 (^644^V) in frame such that the diverged RING domain of Tey is fused C-terminally to the mutant Tey::sfGFP fusion protein, which we therefore name Tey^ΔRNF.sfGFP^ (Fig. 3; Fig. S5).

### Fluorescent antibody staining

Embryo fixation and staining procedures were done as described in (Knirr et al., 1999), larval filets as described in (Fly, 2015), and larval as well as adult midguts as described in (Reim et al., 2012). Filets and midguts were fixed with 3.7% formaldehyde in PBS for 15 min. and after three PBS washes blocked and stained as for the embryos. The following primary antibodies were used: Guinea pig anti-Tey (1:800 (Inaki et al., 2010), visualized with VectaStain Elite ABC kit, Vector Laboratories, and tyramide, PerkinElmer); rabbit anti-RFP (1:250, Rockland); goat anti-GFP (1:500, Genetex); mouse anti-GFP (12A6, 1:100, gift from A. Schambony); chicken anti-βGal (1:200, Abcam); mouse anti-Fas3 (1:30, DSHB, U. Iowa); mouse anti-lamin (T40, 1:30, (Frasch et al., 1988)); mouse anti-integrin αPS2 (1:30, CF.2C7, DSHB, U. Iowa); rabbit anti-β3Tubulin (1:1000, gift from Renate Renkawitz-Pohl); rat anti-Tropomyosin I (1:200, MAC141, Abcam).

Secondary antibodies were generally diluted at 1:200 and included the following: Alexa Fluor 488 conjugated donkey anti-goat (Abcam); Alexa Fluor 555 conjugated donkey anti-rabbit (Abcam); Alexa Fluor 488 conjugated donkey anti-goat (Abcam); Alexa Fluor 488 conjugated goat anti-rabbit (ThermoFisher), DyLight 488 and DyLight 549 conjugated goat anti-rabbit (Jackson ImmunoResearch); Alexa Fluor 555 conjugated donkey anti-rabbit (Abcam); Cy3 conjugated donkey anti-guinea pig (Jackson ImmunoResearch); Alexa Fluor 488 and Alexa Fluor 647 conjugated goat anti-chicken (Abcam); biotinylated goat anti-guinea pig and goat anti-mouse (both 1:500; Jackson ImmunoResearch). Counter-stainings were with Hoechst (Sigma-Aldrich); Alexa Fluor™ *555* and Alexa Fluor™ *647* Phalloidin (ThermoFisher).

### Live imaging

Embryo mountings for GFP/RFP live fluorescent analysis were performed as in (Hollfelder et al., 2014).

### Microscopy

Confocal Z-stacks of fixed specimen were acquired with a Leica SP5 II (20x/0.7 HC PL APO Glycerol, 63x/1.3 HC PL APO Glycerol). Projections of the Z-stacks were performed with FIJI/ImageJ (v1.52v) (Schindelin et al., 2012).

## Results

### Longitudinal visceral muscle precursors actively migrate dorsoventrally and align in parallel

Dorsal and ventral migration of the longitudinal visceral muscle precursors (LVMp’s) takes place in the second half of embryogenesis, in concert with the expansion of the circular visceral muscle precursors (CiVMp’s) along the dorsoventral axis. In order to examine this process more closely, we marked the LVMp’s with *HLH54Fb*-cytoRFP (Fig. 1; depicted in white), the cell surface of the CiVMp’s with anti-Fasciclin3 (Fas3; depicted in magenta), and the nuclei of the CiVMp’s with *bap3*-nGFP (depicted in green). At stage 13, when the cells of the caudal visceral mesoderm (CVM) have completed their anterior migration, the LVMp’s derived from them (after 1 – 2 rounds of myoblast fusion) are spread out along the dorsal and ventral margins, respectively, of the narrow band of CiVMp’s (Fig. 1A, A’; the binuclear CiVMp’s). At early stage 14, when the CiVMp’s elongate dorsoventrally, the LVMp’s elongate in anteroposterior directions and, at the same time, distribute dorsally and ventrally to cover most of the expanded CiVMp’s, except for a central gap (Fig. 1B, B’). This process continues during the remainder of stage 14, when the central gap begins be covered with LVMp’s as well. Importantly, the elongated LVMp’s largely arrange in parallel to their neighbors, although a few of them still tend to be askew (Fig. 1C, C’). During stage 15, upon the dorsal and ventral closure of the midgut, the LVMp’s have evenly dispersed over the midgut along with the extended CiVMp’s. As seen at early stage 16, the elongated LVMp’s have now aligned with their anterior and posterior neighbors and contacted them to form long rows along the length of the midgut, with their dorsal and ventral neighbors being aligned in parallel to them (Fig. 1D, D’). During stage 16, this arrangement matures such that the differentiating longitudinal gut muscles form thin muscle fibers, each one extending over much of the length of the midgut, that are distributed roughly equidistantly around the midgut and orthogonally to the circular visceral muscles (Fig. 1E, E’).

**Fig. 1.**
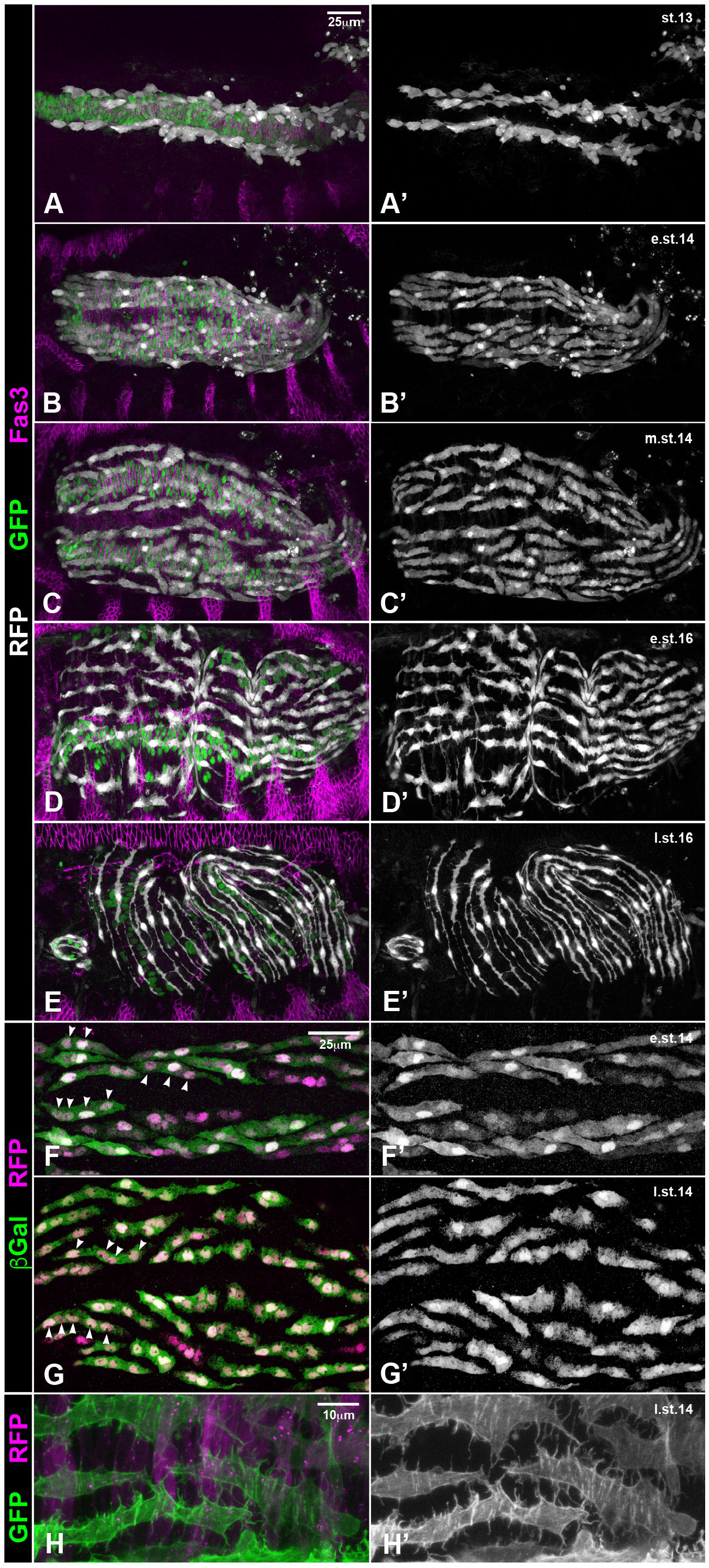
Dorsal and ventral migration of longitudinal visceral muscle precursors. Left hand column shows composite images and right hand column depicts channels showing visceral muscle (LVMu) precursors only. (A - E’) Unless noted otherwise, anterior is to the left and dorsal is up in this and all other figures. Shown is the migration of longitudinal visceral muscle precursors in fixed embryos of consecutive stages that express RFP in the LVMu precursors (from *HLH54Fb-cytoRFP*; white) and nuclear GFP in circular visceral muscle (CiVMu) precursors (from *bap3-nGFP*; green). The CiVMu cell outlines are visualized by Fasciclin 3 (Fas 3) stainings (magenta), which are decreasing in the visceral mesoderm after stage 14 while increasing in the epidermis. Scale bar in (A) applies to (A – E’). (F – G’) Nuclear labeling for Histone-RFP shows that LVMu’s migrate as syncytia containing 3 – 5 nuclei from early stage 14 onwards (arrow heads; genotype: *UAS-His-RFP/+;tey^5053A^ UAS-lacZ/+*). Scale bar in (F) applies to (F – G’). (H, H’) Direct imaging of heat-fixed embryos (genotype: *UAS-Lifeact-GFP/bap3-RFP; tey^5053A^/TM6 Dfd-EYFP*) shows that the LVMp’s (green) migrate and align on top of the CiVMu precursors (magenta) in the presence of pronounced filopodia that extend towards their neighbors.

The LVMp’s undertake their dorsoventral migrations as syncytia, which we examined by driving (largely) cytoplasmatic LacZ together with nuclear histone-RFP in them. At the beginning of this migration process during early stage 14, most of the migrating LVMp’s already contain two to four nuclei (Fig. 1F, F’), during late stage 14 three to five nuclei (Fig. 1G, G’), and at the end of migration, before lining up at stage 15, most of them have acquired 6 nuclei (data not shown).

Of note, the migrating LVMp syncytia feature prominent filopodia distributed over their entire cell surface. In addition to contacting the underlying CiVMp’s as their migration substrate, many of these filopodia extend laterally and contact filopodia or cell bodies of neighboring LVMp’s (Fig. 1H, H’; see also Fig. 1C’, D’). Time lapse videos show that these filopodia are highly dynamic, presumably in sensing the migration substrate and the presence of neighboring cells (Movie 1; Movie S1). These filopodia still show dynamic activity during the parallel arrangements of differentiating LVMu’s during stage 16 and only start disappearing at the end of this morphogenetic process at late stage 16 (Fig. 1E’; Movie S1).

### Tey is expressed in the caudal visceral mesoderm and the migrating LVMp’s, as well as in founders of somatic muscle M12

Previously, we presented the expression of Tey in differentiated neurons and somatic muscle M12 (Inaki et al., 2010). Herein, we describe Tey expression during the migration of visceral and somatic mesodermal cells. To put the Tey-stained cells into tissue context we performed these stainings in the presence of *Fas3-nGFP*, which marks the trunk visceral mesoderm (TVM) and the circular visceral muscles (CiVMu’s) derived from it. Tey is already expressed in the CVM prior to its anterior migration (data not shown) and at stage 11 is detected in the bilateral clusters of CVM cells that have migrated towards the posterior-most clusters of the TVM (Fig. 2A; see also Bae et al., 2017). Tey expression in CVM cells and their descendent LVMp’s persists during their anterior and subsequent dorsoventral migration (Fig. 2B, C) and is still present in the differentiating LVMu’s at stage 16 (Fig. 2D). In addition, at stage 12 – 13 Tey expression initiates in a single cell per hemisegment in the somatic mesoderm, which corresponds to the founder of muscle M12 (Fig. 2B). During myoblast fusion at stage 14, Tey expression is still prominent in the M12 precursors but decreases during stage 15 and thereafter (Fig. 2C, D). As already discernible in Fig. 2A – D, high magnification views of migrating CVM cells co-stained for Tey, nuclear envelopes, and DNA confirmed that the Tey protein is located strictly in the nuclei of these cells (Fig. 2E, E’).

**Fig. 2.**
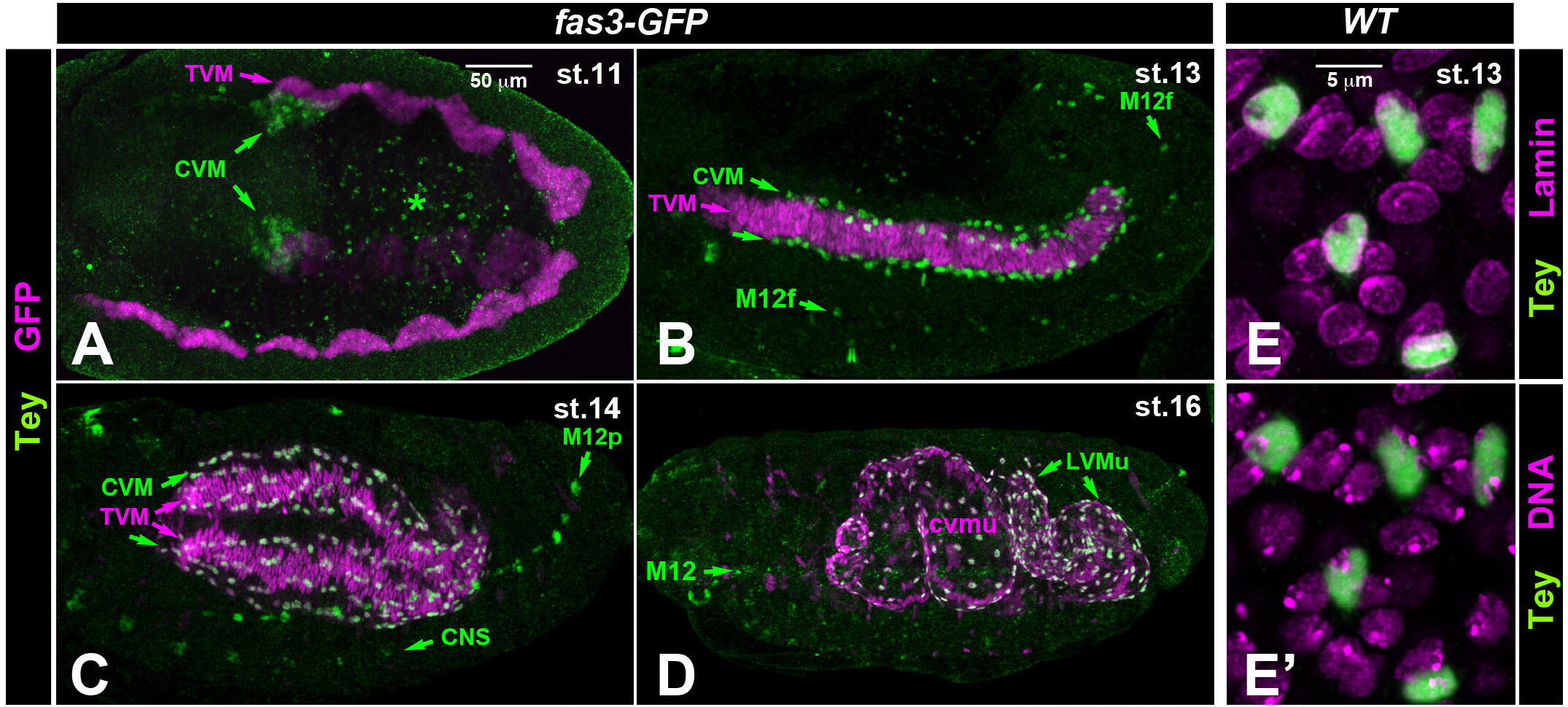
Expression and nuclear localization of the Tey protein. (A – D) anti-Tey (green) + anti-GFP (magenta) - stained *Fas3-nGFP* embryos (see scale bar in A). (A) At stage 11, Tey is detected in the bilateral caudal visceral mesoderm (CVM) cell clusters that have migrated towards the posterior-most cluster of the trunk visceral mesoderm (TVM). (B) At stage 13, Tey is present in the nuclei of the CVM cells that have migrated anteriorly along the dorsal and ventral margins of the TVM, as well as in the nuclei of single somatic muscle founder cells, M12f, within each segment. (C) At stage 14, nuclear Tey is present in the LVMp’s migrating along the TVM, as well as in muscle 12 precursors (M12p‘s). CNS expression also appears. (D) At stage 16, nuclear Tey is present in all longitudinal visceral muscles (LVMu’s), but is diminished in somatic M12. (E, E’) High magnification views confirming nuclear localization of Tey. (E) Anti-Tey + anti-Lamin staining and (E’) anti-Tey + Hoechst (DNA) staining from the same stage 13 embryo (see scale bar in (A)). Tey is seen strictly within the confines of the nuclear envelope and within the area of the DNA.

### Tey is a diverged member of the RNF220 family of proteins

Sequence alignments and comparisons show that Tey is a member of the RNF220 protein family, which has representatives in all well-characterized vertebrate and most arthropod species. These proteins share a highly conserved domain in their center, termed RNF220 domain, as well as a C-terminal RING finger domain (Fig. 3, Fig. S1). Various other stretches, such as the N-terminus of Tey, are conserved in the orthologs of other species as well (Fig. 3, Fig. S1). Of note, the RING finger domain of Tey is highly diverged when compared to the those of its orthologs in other dipterans, coleopterans, and vertebrates, and only contains a partial RING finger motif (Fig. S1). Hence, it is questionable whether the C-terminal domain of Tey is able to mediate ubiquitin ligase activity, as has been reported for mouse RNF220 (Ma et al., 2019; Song et al., 2020; Ma and Mao, 2022). However, *Drosophila* has a paralog of Tey of largely unknown function, CG4813 (recently renamed *Drosophila* RNF220; Sferra et al., 2021), which features a complete RING domain in addition to its RNF220 domain, and in *Tenebrio*, both paralogs have a complete RING domain (Fig. 2, Fig. S1). The tertiary structures of *Drosophila* Tey and mouse RNF220 have been predicted by AlphaFold (Senior et al., 2020). As might be expected from the sequence alignments, the RNF220 domains are predicted to have clear structural similarities, whereas at their C-termini no clear similarities are evident (Fig. S1).

**Fig. 3.**
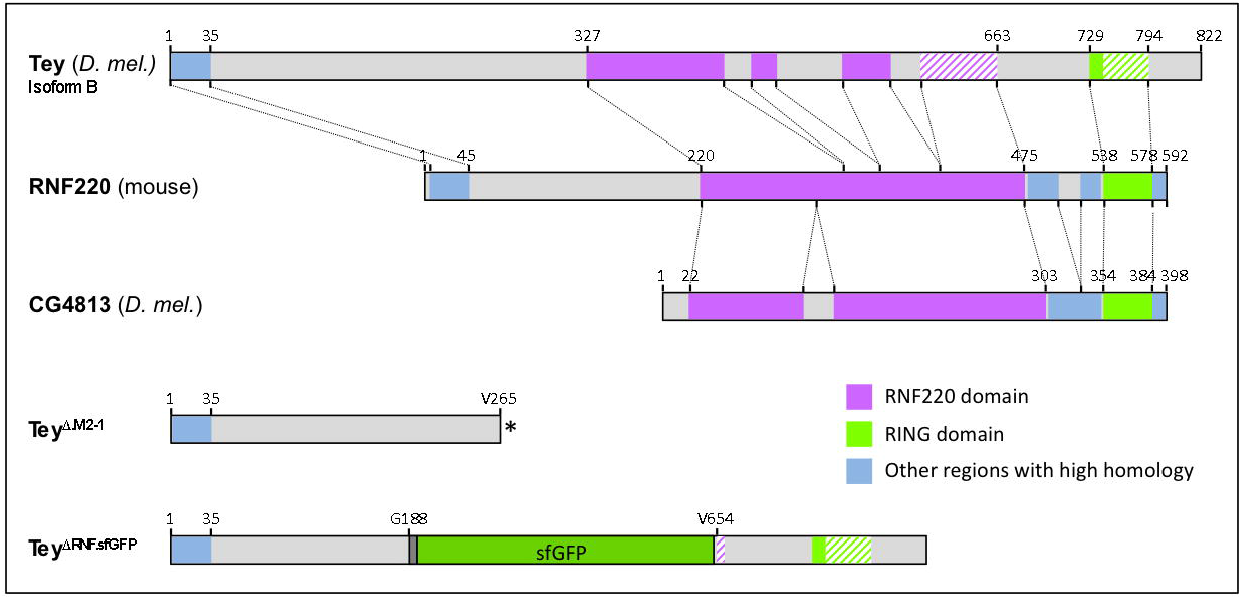
Structure and conservation of wild type and mutated versions of Tey and related proteins. (Top) Shown are the schematic alignments of highly conserved domains between Tey (*D. mel*.), RNF220 (*M. musculus*), and CG4813p (*D. mel*.). (Bottom) Schematics of the predicted Tey and Tey::sfGFP (superfolderGFP) fusion proteins translated from the respective CRISPR/Cas9-derived *tey* alleles.

### *tey* is required for normal dorsoventral migration, including the attainment of proper orientation and spacing, of LVMu’s

In addition to the line *tey^5053A^*, in which a Gal4 insertion at the 5’ end of *tey* leads to undetectable expression levels of *tey* mRNA (Inaki et al., 2010) and protein (Fig. S2), we generated additional alleles via CRISPR/Cas9 to increase confidence that we observe null phenotypes and to exclude possible effects of closely linked second site mutations. In two of these alleles, *tey^Δ.M2-1^* and *tey^Δ.M1-11^*, deletions from exon 4 to exon 8 cause the predicted native ORF to terminate at V^265^ and G^266^, respectively, which leads to a complete absence of the RNF220 and the diverged RING finger domains (Fig. 3, Fig. S1). A third allele, *tey^ΔRNF.sfGFP^*, creates a version of the Tey protein in which a large central portion that includes the RNF domain is replaced with sfGFP as a tag (Fig. 3, Fig. S1; see Materials & Methods).

Until early stage 14, when the anteroposterior migration of the CVM has been completed and the LVMp’s have started forming elongated syncytia that begin to disperse, we do not observe any abnormalities in *tey* mutants (Fig. 4A, A’, cf. Fig. 1B, B’). However, further progression of their dorsoventral migration during mid and late stage 14 is highly aberrant in embryos lacking *tey* activity. Rather than being aligned quite faithfully in anteroposterior orientations and being distributed evenly along the dorsoventral axis, in *tey* mutant embryos the LVMp’s frequently assume oblique or even transverse orientations, and their distances from one another are uneven. Furthermore, the shape of the syncytia becomes variable and abnormal during this stage in that they tend to assume more squat or triangular shapes instead of the elongated spindle-like shapes seen in the controls. (Fig. 4B – C’, arrow heads, cf. Fig. 1C, C’; Fig. S3B, cf. Fig. S3A). Their nuclei tend to be arranged in clusters instead of in single anteroposterior files within the LVMp syncytia (e.g., Fig. 4C, C’, cf. Fig. S3A). At stage 15 and early stage 16, these defects in LVMp migration and morphogenesis become even more pronounced. Although LVMp migration does reach the dorsal and ventral areas of the forming midgut tube, in contrast to their normal behavior this migration appears very uncoordinated because the LVMp’s are oriented in seemingly random directions. Additionally, while some LVMp’s are crowded together, other areas of the midgut remain devoid of them (Fig. 4D – E’, cf. Fig. 1D, D’; Fig. S3D, cf. Fig. S3C; Fig. S3F, cf. Fig. S3E; Fig. S3N, cf. Fig. S3M). Our finding that there is no increased apoptosis in *tey* mutant LVMp’s suggests that the observed empty areas are a direct result of aberrant migrations rather than cell death in the absence of *tey* function (Fig. S3L, cf. Fig. S3K). During stage 16, some of the LVMu’s do manage to extend in anteroposterior directions along the gut tube, but even these maintain abnormal shapes and the coverage of the midgut by LVMu’s remain very uneven (Fig. 4E – G, cf. Fig. 1E, E’; Fig. S3D, cf. Fig. S3C; Fig. S3H, cf. Fig. S3G, Fig. S3J, cf. Fig. S3I). Time lapse videos further illustrate the migration and morphogenesis defects of LVMp’s in the absence of *tey* function that we described above from static pictures, particularly the abnormally close contacts that are frequent among migrating LVMp’s (Movie 2) and the resulting disorganized arrangements and morphologies of differentiated LVMu’s at late stage 16 (Movie S2). Filopodial extensions are still present but we have not quantified their dynamic properties in comparison to wildtype controls.

**Fig. 4.**
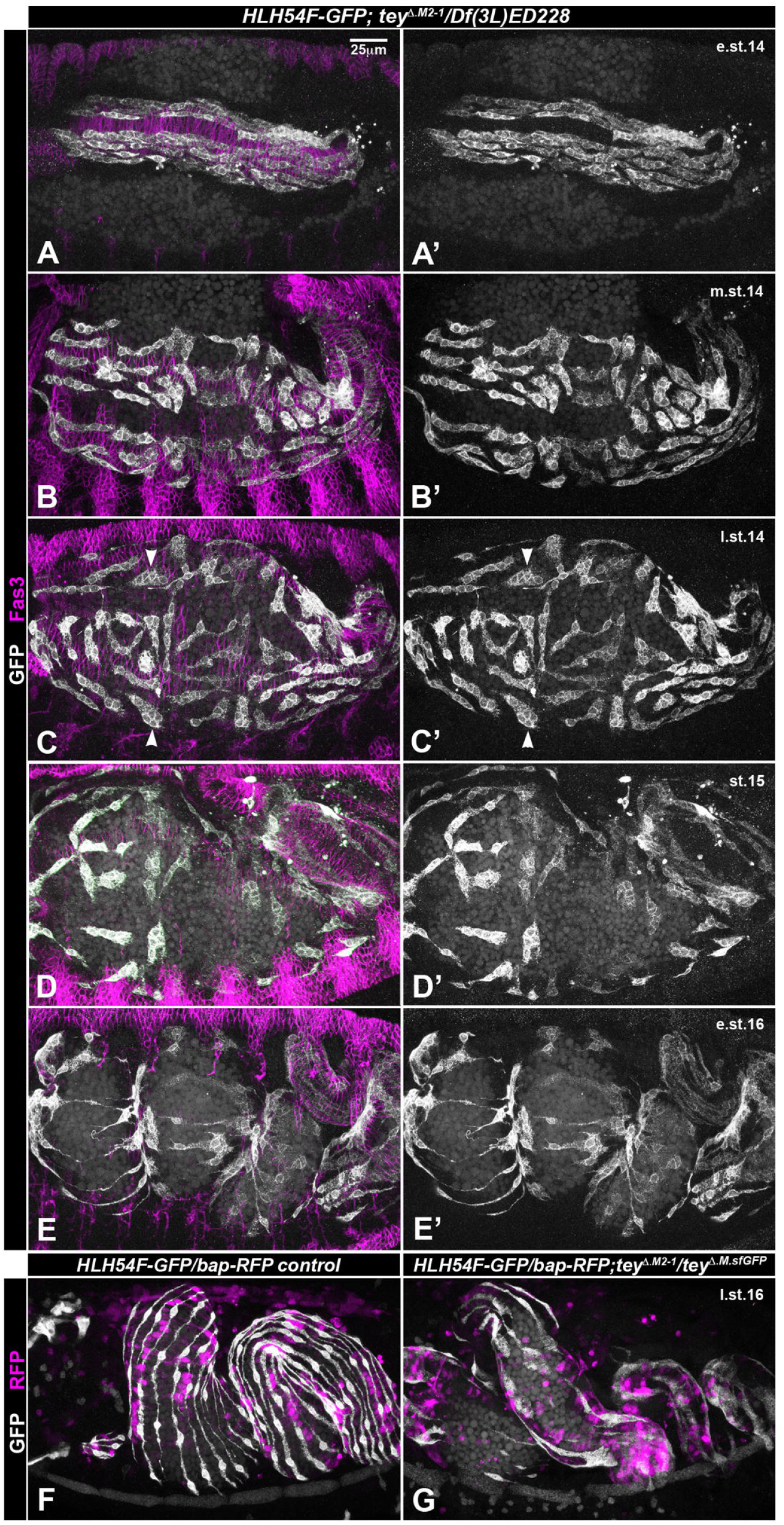
Mis-migration of longitudinal visceral muscle (LVMu) precursors in *tey* mutant embryos. (A – E’) Anti-GFP (white) + anti-Fas3 (magenta)-stained *tey* mutant embryos with the genotype shown on the top of the panel (left column: dual channels; right column: single GFP channel; see (A) for scale bar). See Fig. 1 for comparisons with the wildtype situation. (A, A’) At early stage 14, anterior migration of the LVMp’s and initiation of their dorsal and ventral migration is not visibly affected in the absence of *tey* function. (B – C’) At mid and late stage 14, an increasing proportion of the LVMp’s is oriented obliquely or transversely instead of longitudinally in parallel, and display abnormal shapes. (D, D’) At stage 15, the LVMp’s have completed their migration around the midgut but show random orientations and shorter as well as broader shapes. Some midgut areas are devoid of LVMp’s. (E) At stage 16, the LVMu’s partially attain longitudinal orientations but with irregular cell shapes and distances from one another. The main cell bodies tend to occupy the gut constrictions. (F) Stage 16 control embryo (either *tey^Δ.M2-1^* or *tey^ΔRNF.sfGFP^/TM6 Dfd-EYFP*) carrying *HLH54Fb-cytoRFP/bap3-RFP* on the 2^nd^ chrom. and stained for GFP (white) in the LVMu’s and for RFP in the CiVMu’s. The LVMu’s are oriented roughly equidistantly and in parallel along the length of the midgut. (G) Stage 16 *tey^Δ.M2-1^/tey^ΔRNF.sfGFP^* embryo with reporters and staining as in (F). The LVMu’s are not strictly oriented in anterior-posterior directions along the midgut, display variable abnormal shapes, and frequently contact their neighbors. The midgut also has areas lacking LVMu’s.

As a result of the seemingly random migration events in mutant embryos, the final LVMP and LVMu arrangements are quite variable in individual embryos, but all allelic combinations tested show essentially the same range of defects and thus we believe that all of the presented alleles are null. Anti-RFP stainings of the truncated Tey::sfGFP fusion protein in *tey^ΔRNF.sfGFP^* embryos shows that the residual Tey peptide is cytoplasmic instead of nuclear (Fig. S3M – N’) and the same presumably is the case for Tey^Δ.M2-1^ and Tey^Δ.M1-11^ (if stable).

### Tey is required for proper arrangement, morphogenesis, and differentiation of LVMu’s

The viability of *tey* mutant larvae (albeit decreasing until 3^rd^ instar) and the survival of a low percentage of adult escapers allowed us to examine the role of *tey* in the morphogenesis and differentiation of LVMu’s. Continued Tey expression in LVMu’s until at least 3^rd^ instar (Fig. 5E) indicates that *tey* may still be required for the continued differentiation and homeostasis of these muscles growing during these stages. Indeed, compared to the almost equidistantly spaced, thin LVMu’s aligned in parallel (Fig. 5A, C), the LVMu’s in *tey* mutant larval midguts exhibit highly unordered arrangements and aberrant shapes. Although they mostly tend to be oriented in anteroposterior directions, some of them are oriented obliquely or are curved instead of being straight. The mutant larval LVMu’s are not aligned into long fibers in anteroposterior directions, show highly variable distances to their lateral neighbors, and often touch or even cross them (Fig. 5B, B’, D, D’, F, F’, H, H’; cf. 5A, C, G, respectively). In part, these phenotypes may be due to the above-described migration defects during embryonic development. However an additional phenotype in 3^rd^ instar larvae concerns the morphology of the mutant LVMu’s and the arrangement of their myofibrils. Specifically, the mutant LVMu’s are much broader, sometimes are branched or split at their ends, and their actomyosin fibrils are spaced apart along their lengths and often display frayed arrangements (Fig. 5B, B’, D, D’, F, F’, H, H’; cf. 5A, C, G; Fig. S4B, cf. Fig. S4A). Hence, *tey* apparently continues to be required for proper morphogenesis and differentiation of the LVMu’s during the final embryonic and the larval stages. Nevertheless, striations are formed and show the same range of spacing as in control larvae (e.g., Fig. S4D, cf. Fig. S4C).

**Fig. 5.**
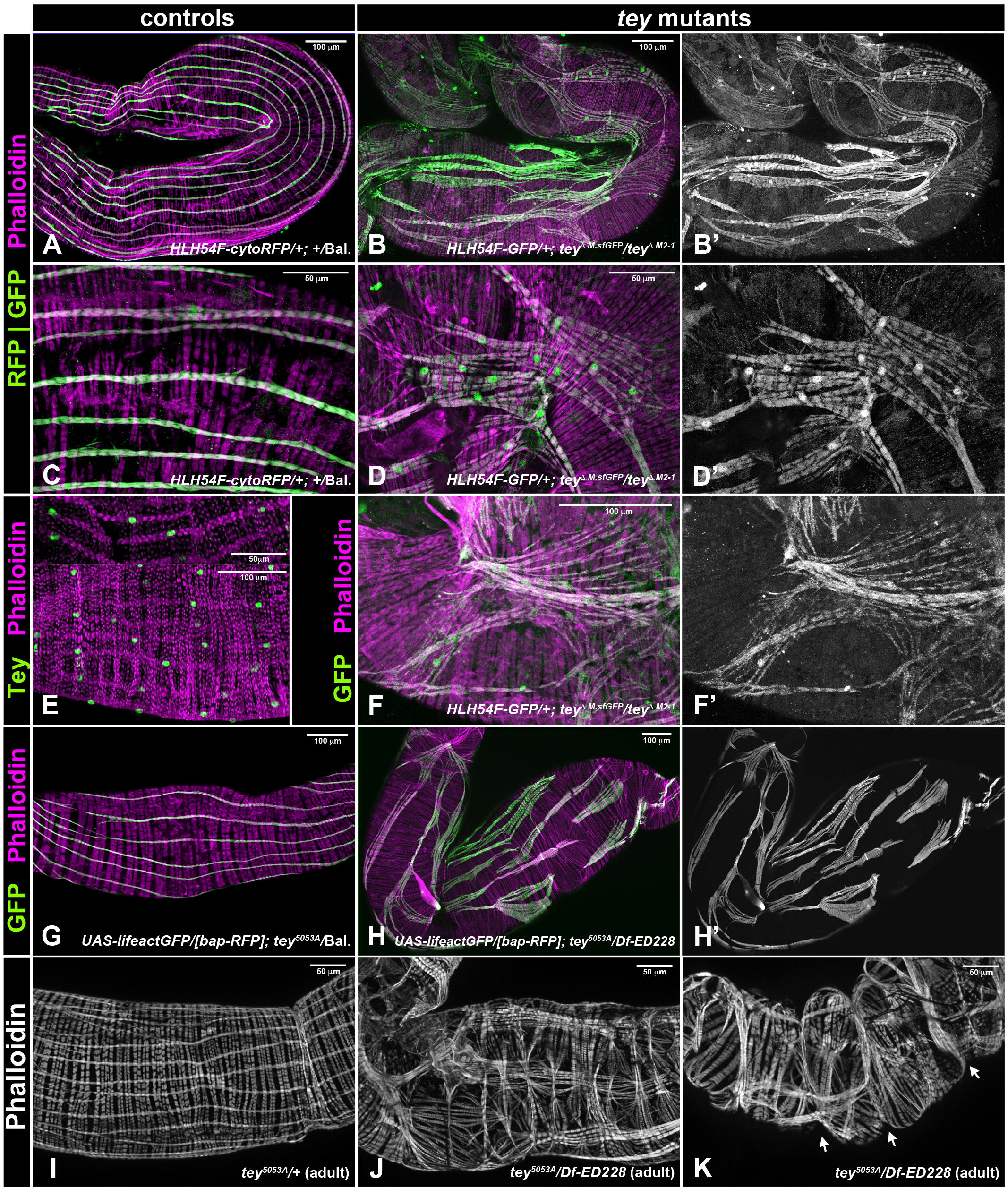
Abnormal morphologies and differentiation of longitudinal midgut muscles in *tey* mutant 3^rd^ instar larvae and adults. (A, C) 3^rd^ instar larval midguts from control animals (*+/TM6-Dfd-EYFP*) carrying *HLH54Fb-cytoRFP* as an LVMu marker, shown at two different magnifications (see scale bars) and stained for RFP (depicted green) and F-actin (magenta). The LVMu’s form slender muscles that are oriented in parallel and roughly equidistantly around the midgut. (B, B’, D, D’, F, F’) 3^rd^ instar larval midguts from *tey* mutant animals (*tey^ΔRNF.sfGFP^/tey^Δ.M2-1^*) carrying *HLH54Fb-GFP* as an LVMu marker, shown at three different magnifications (see scale bars) and stained for GFP (green) and F-actin (magenta). The LVMu’s display severe abnormalities with respect to their shapes, orientations, and myofibril alignments. The LVMu’s often touch their neighbors and are not strictly arranged in anterior/posterior orientations along the midgut. (E) Larval 3^rd^ instar midguts stained with anti-Tey (green) and fluorescent phalloidin (magenta), showing continued expression of nuclear Tey in larval LVMu’s. (G, H, H’) 3^rd^ instar larval and F-actin (magenta) (anti-RFP channel not shown due to the disappearance of *bap*-driven RFP in larvae). (H, H’) Larval midgut from *tey^5053A^/Df(3L)ED228* animal with reporters and staining as in (G), showing similar LVMu abnormalities as with the allele combination depicted above. (I) Adult control midgut (from *tey^5053A^/+*) stained for F-actin, showing regular spacing and orthogonal arrangements of midgut muscles as in larvae. (J, K) *tey* mutant midguts from adult escapers of the genotype *tey^5053A^/Df(3L)ED228*, showing similar LVMu phenotypes as *tey* mutant larvae. In addition, CiVMu arrangements are locally distorted and often there are midgut constrictions along the A/P axis (K; arrows).

Because the LVMu’s are completely reconstituted from scratch during metamorphosis (Aghajanian et al., 2016) it was interesting to determine whether *tey* is required during this process as well. Indeed, the newly built LVMu’s from adult escaper flies show severe abnormalities, with aberrancies that are similar to the ones in mutant larval midguts. (Fig. 5J, K; cf. Fig. 5I). In addition, these abnormalities indirectly affect the arrangements of the CiVMu’s, which tend to be bundled together in the areas underneath the LVMu’s. This effect may be due to improper attachments between LVMu’s and CiVMu’s, which also could account for the “accordion-like” shape of the midgut that could result from contracting and incorrectly attached LVMu’s in mutant flies (Fig. 5K, cf. Fig. 5I, Fig. S4F, cf. Fig. S4E). In sum, we find that *tey* is required from embryonic stage 14 until early adult stages for the proper migration and differentiation of the LVMu’s and their precursors.

### Tey regulates the migration of somatic muscle 12 precursors

Tey is expressed in the founders and precursors of somatic muscle 12 (M12) (Inaki et al., 2010; Fig. 2B, C, D; Fig. 6A). Normally, M12 muscles (aka, VLM1) are positioned at the dorsal margin of the band of ventral longitudinal muscles (VLMs) and align at their epidermal attachment sites with their anterior and posterior M12 neighbors to form a linear row of M12 muscles along the a/p axis. By contrast, in *tey* mutant embryos each M12 muscle is oriented obliquely within each segment, such that these muscles form a zig-zag pattern along the length of the abdomen (teyrha meyrha, 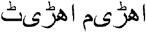, Urdu, = zig-zag). Closer inspection with co-stainings for the other, unaffected VLMs or their epidermal muscle attachment sites shows that the anterior attachment of each M12 usually is shifted ventrally by more than the width of the muscle, whereas the posterior attachment is usually shifted by at least two muscle widths (Fig. 6B, D, F; cf. Fig. 6A, C, E). Thus, in addition to the orientation defects, each M12 is shifted ventrally as a whole.

**Fig. 6.**
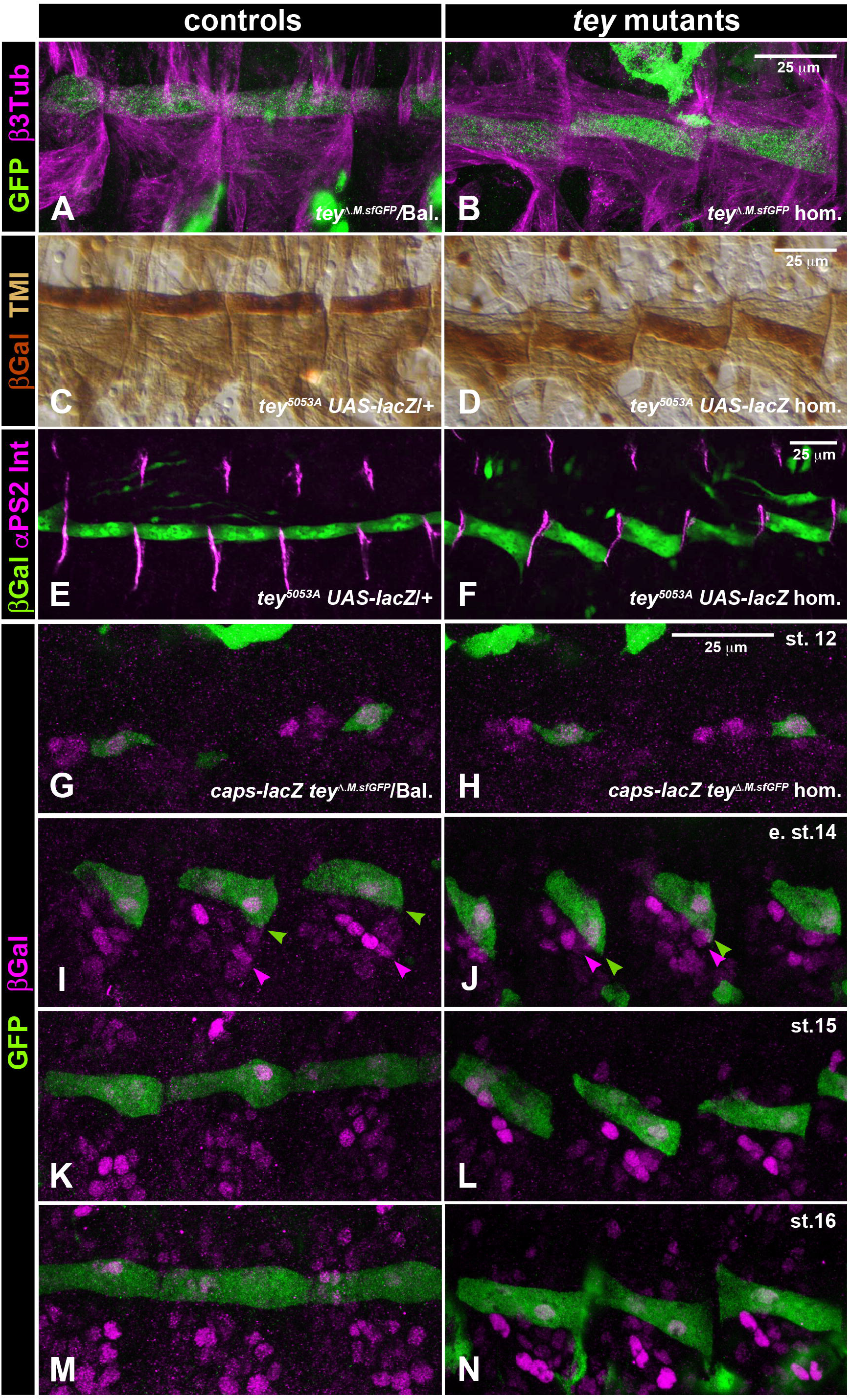
Aberrant position and migration of somatic muscle 12 (M12) and its precursor during embryonic muscle development. Left column shows control embryos and right column shows *tey* mutants with the same reporters, stainings, and magnifications as the corresponding controls (see scale bars in right column; scale bar in (H) applies to (G – N). (A – F) show fully differentiated muscles at stage 16. (A, B) As compared to the control (*tey^ΔRNF.sfGFP^/TM6 Dfd-EYFP*), M12 muscles in a homozygous *tey^ΔRNF.sfGFP^* mutant (stained for TeyΔRNF::GFP in green and β3-Tubulin in magenta) are shifted ventrally and display oblique orientations. (C, D) As with *tey^ΔRNF.sfGFP^* mutants, homozygous *tey^5053A^* embryos (*tey^5053A^ UAS-lacZ*; (D)) display ventrally-shifted M12 muscles with oblique orientations as compared to the controls (*tey^5053A^ UAS-lacZ/+*; (A), both stained for βGal expression in M12 (dark brown) and topomyosin I in all muscles (TM I; light brown)). (E, F) Stainings of control and *tey* mutant embryos, respectively, with the same genotypes as in (C, D) for M12 with anti-βGal + anti-αPS2 integrin highlight the ventral shifts of M12 attachments to the segmental muscle attachment sites in the mutant. (G – N) Developmental series showing the abnormal migration of M12 precursors in tey mutant (*caps-lacZ tey^ΔRNF.sfGFP^* homozygotes) as compared to the controls (*caps-lacZ tey^ΔRNF.sfGFP^/TM6 Dfd EYFP*). M12 precursors are marked with anti-GFP (for TeyΔRNF::sfGFP; green) and anti-βGal (for the nuclear M12 marker *caps*-LacZ, magenta). (G, H) At stage 12, the M12 founder cells feature identical positions and morphologies in control and mutant embryos. (I) In early stage 14 control embryos, the M12 muscle precursors extend towards their future anterior attachment sites within each segment, which involves an arched path of migration. (J) In early stage 14 *tey* mutants, M12 migration occurs in dorsal/anterior directions without arching back ventrally at their anterior ends. In comparison to the controls, the entire cell bodies of M12 precursors have shifted ventrally relative to their unaffected *caps*-LacZ-positive neighbors (see arrow heads). (K) After completing their anterior migration at stage 15, M12p’s in controls establish contacts with their attachment sites in each segment at the same dorsoventral positions as their M12p neighbors. (L) In stage 15 *tey* mutant, the bodies of the M12p’s remain shifted ventrally, with their posterior attachments showing a stronger ventral shift as compared to their anterior attachments. (M) After final differentiation and maturation of their attachments at stage 16, M12 muscles in the control form a continuous band of longitudinal muscles in abdominal segments. (N) In a stage 16 *tey* mutant embryo, M12 muscles are arranged as a zig-zag.

To define the specific role of *tey* in M12 founder and precursor migration, we compared a developmental series of heterozygous and homozygous *tey^ΔRNF.sfGFP^* embryos that were co-labeled for Tey::sfGFP and the reporter *caps*-LacZ. The latter (nuclear) marker is co-expressed with Tey in founders and precursors of M12, as well as in several unaffected neighboring muscle founders and precursors that can serve as positional landmarks. As seen in Fig. 6G, H, the Tey::sfGFP^+^ *caps*-LacZ^+^ M12 founders are shaped normally and are located at the same positions with respect to their anterior Tey::sfGFP^-^ *caps*-LacZ^+^ neighbors in both control and *tey* mutant embryos. In addition, *caps*-LacZ expression in the mutant founder cells shows that they maintain M12 founder cell identities in the absence of *tey* function. Upon myoblast fusion at early stage 14, it is largely the anterior end of each nascent M12 myotube that migrates towards its anterior epidermal attachment site, whereas the posterior end is already positioned close to the posterior attachment site to which it attaches during mid stage 14 (Fig. 6I). Normally, the leading edges of migration of M12p’s initially migrate in a dorso/anterior direction. From mid stage 14, they slightly bend down to contact their anterior epidermal attachment sites at the position where the posterior attachment of their anterior M12 neighbor is attached, thus eventually forming the linear arrangement of M12 muscles (Fig. 6I, K, M). In the absence of *tey* function, the leading edges of M12p’s also migrate in a dorso/anterior direction. However, when compared to the neighboring *caps*-LacZ^+^ muscle precursors, it becomes clear that the entire cell bodies of M12 are positioned more ventrally, and particularly so at their posterior sides, in *tey* mutants as compared to the controls (Fig. 6J, cf. Fig. 6I; see arrow heads). During stage 14, these muscle precursors retain their incorrect orientations and positions and during stages 15 – 16 establish their abnormal anterior and posterior attachment sites, which then leads to the observed zig-zag pattern of M12 muscles (Fig. 6L, N; cf. Fig. 6K, M). We favor the interpretation that normally M12 founders and precursors migrate a short distance dorsally from their place of origin, where their leading edges can respond to pathfinding signals from their native attachment sites. However, in the absence of *tey* function they fail to undergo this dorsal migration and thus remain in the vicinity of the Tey^-^ Caps^+^ precursors, where they erroneously respond to pathfinding cues for their epidermal attachment from tendon cells that are closer to their more ventral position.

Additional information on the mis-positioning of M12 in *tey* mutants can be gleaned from the analysis of 3^rd^ instar larval muscle patterns. As shown in Fig. 7B, D, F, the mutant M12 muscles are clearly shifted ventrally. Their oblique orientation is a result of the attachments at the anterior segment border to positions overlapping with the attachments of M6 (aka, VLM3) and their posterior attachments are shared with those of M7 (VLM4) (cf. Fig. 7A, C, E). Additionally, many mutant larvae show examples of M12 muscles that display split (Fig. 7D) or broadened (Fig. 7F) attachments, indicating that muscle attachments at their more ventral positions cannot be established as consistently as at their normal positions. Similar to the gut muscle phenotypes, the observed somatic muscle phenotypes are identical for all allelic combinations tested (Fig. 7).

**Fig. 7.**
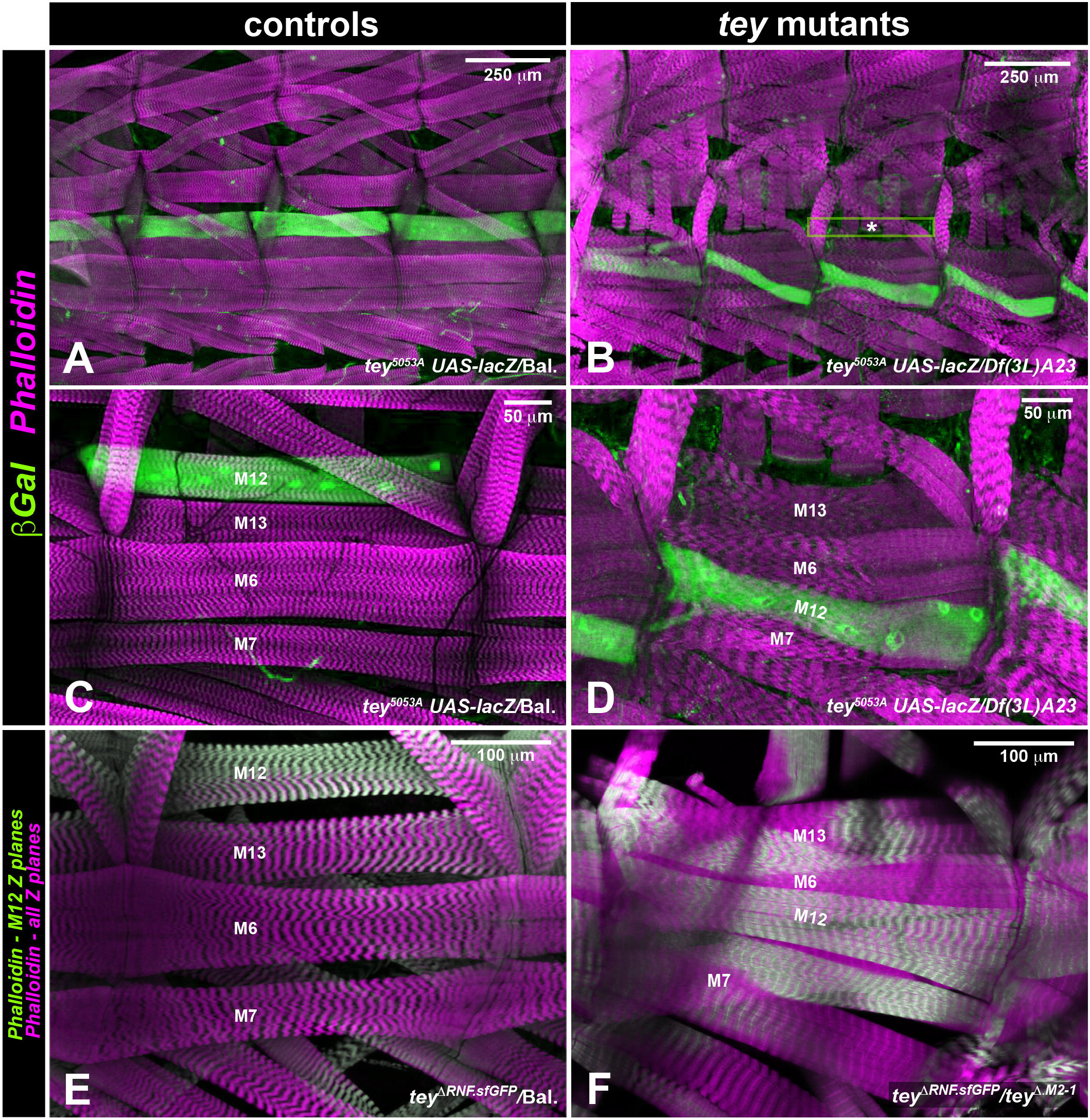
Phenotypes of M12 muscles in *tey* mutant 3^rd^ instar larvae. Left hand column shows controls and right hand column shows mutants stained for the same markers. Animal in (B, D) was at mid 3^rd^ instar and all others at late 3^rd^ instar. (A – D) Whereas in control larvae (*tey^5053A^ UAS-lacZ/TM6 Dfd-EYFP*) the M12 muscles form a longitudinal muscle band at the dorsal-most position along the ventral longitudinal muscles, in a *tey* mutant larva (*tey^5053A^ UAS-lacZ/Df(3L)A23*) M12 muscles form a zig-zag pattern. (E, F) Phalloidin-stained ventrolateral muscles of *tey* mutant 3^rd^ instar larva (*tey^ΔRNF.sfGFP^/tey^Δ.M2-1^*) vs. control larva (*tey^ΔRNF.sfGFP^/TM6 Dfd-EYFP*). For clarity, the confocal Z planes that included M12 are rendered in green, whereas those containing the other muscles are rendered in magenta. Comparison of M12 position and orientation in (F) vs. (E) shows that this allelic combination provokes the same M12 muscle phenotype as the one shown above.

### Overexpression and ectopic expression disturbs muscle precursor migration and morphogenesis

Forced overexpression of *tey* in the LVMp’s via *tey-GAL4* (aka, *tey^5053A^*) and *UAS-tey* in the presence of *UAS-lacZ* as a marker showed that an excessive dosage of *tey* is detrimental to the proper migration and morphogenesis of LVMp’s. As seen in Fig. 8B (cf. Fig. 8A), the LVMp’s appear to migrate out prematurely from the dorsal and ventral cell rows to cover the band of CiVMp’s and do not display regular spindle shapes oriented along the a/p axis in this genetic background. At late stage 14, the LVMp’s with *tey* overexpression likewise show irregular orientations and shapes, as well as an uneven coverage of the midgut (Fig. 8D, cf. Fig. 8A).

**Fig. 8.**
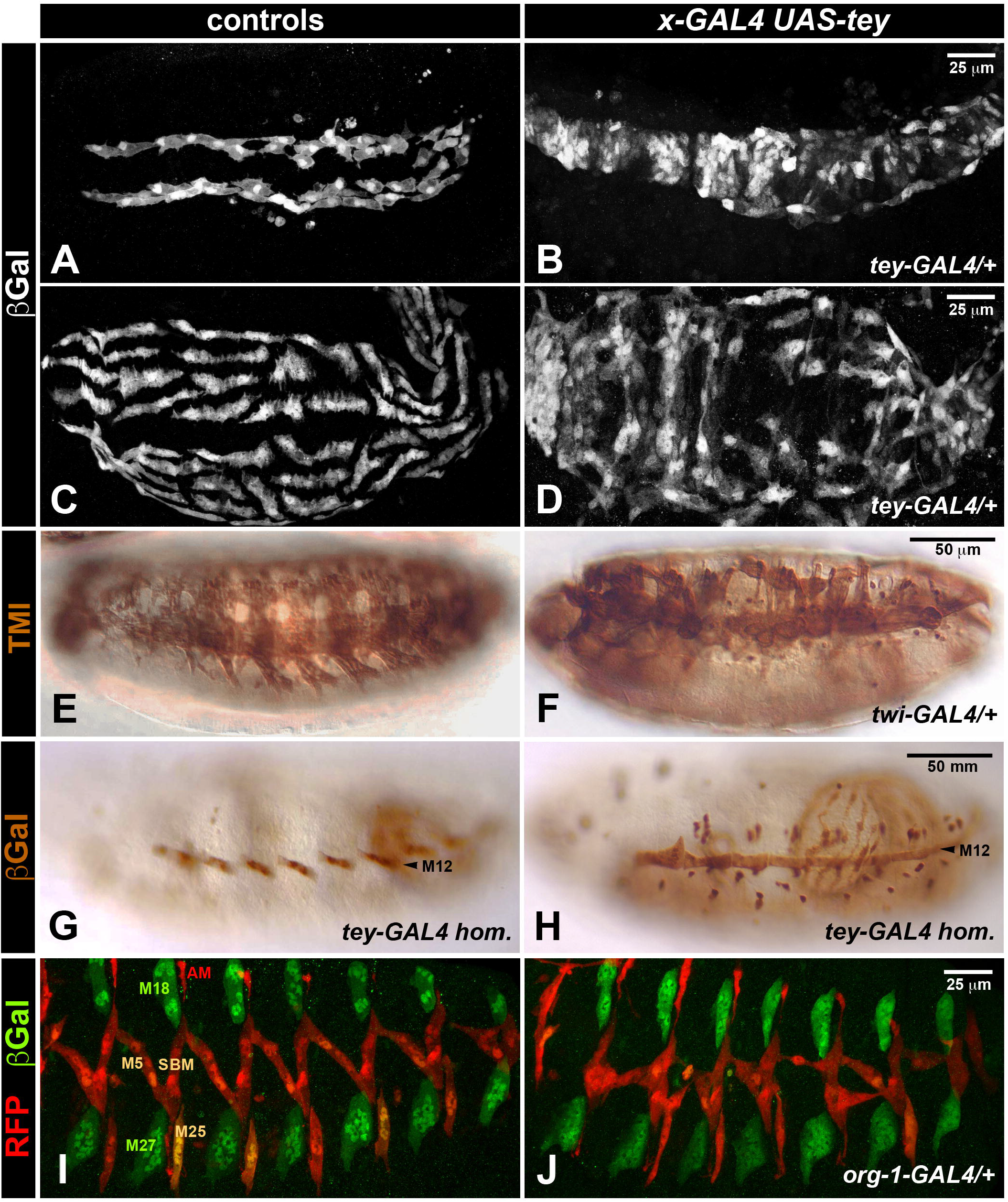
Disrupted muscle migration and morphogenesis upon forced *tey* expression in developing longitudinal visceral muscles and somatic muscles. Left hand column shows control embryos and right hand columns embryos containing both *UAS-tey* and various GAL4 drivers (denoted as *X-GAL4*, where the genotypes of X are provided on the respective panels). (A, B) As compared to the control (*HLH54Fb-lacZ UAS-tey*, stained for βGal (A)), in stage 13 embryo with *tey* overexpression within LVMp’s via *tey-GAL4* (from *tey^5053A^*) these cells prematurely spread over the entire width of the TVM. (C, D) Genotypes and staining as in (A, B). As compared to the control, stage 14 embryo with overexpression of *tey* shows abnormal orientations and shapes of LVMp’s. (E, F) Somatic muscle patterns in stage 16 embryos visualized with anti-tropomyosin I show severely disrupted muscle morphologies when *twi-GAL4* is used to drive ectopic *tey* expression in the mesoderm. (G, H) Forced expression of *tey* in a homozygous *tey* mutant background (H; *UAS-tey/+; tey^5053A^ UAS-lacZ* hom.) rescues the zig-zag pattern seen in *tey* mutants (G; *tey^5053A^ UAS-lacZ* hom.) and produces almost normal M12 morphologies. (I) Stage 16 control embryo (*RRHS59-lacZ/+; HN39org-1-GAL4 S18org-1-RFP/+*) showing the normal somatic muscle patterns of the *org-1* expressing muscles M5, M25, SBM and alary muscles (AM) (anti-RFP, red) and the *slou* expressing muscles M5, M25, SBM, M18, M27 (anti-βGal, green). (J) In stage 16 embryo with ectopic expression of *tey* via *org-1-GAL4 (RRHS59-lacZ UAS-tey/+; HN39org-1-GAL4 S18org-1-RFP/+*), the muscles with ectopic *tey* (particularly M5, M25, and SBM) display severely abnormal orientations and shapes. By contrast, the *slou*-specific muscles M18 and M27 lacking GAL4 activity are normal and can serve as landmarks.

Because in the somatic mesoderm *tey* normally is only expressed in a single muscle precursor, M12p, among the 30 muscle precursors within each hemisegment, we tested whether the presence of *tey* would alter the development of muscle precursors in which it is not normally expressed. As shown in Fig. 8F, ectopic expression of *tey* with an early pan-mesodermal driver, *twi-GAL4*, leads to severe disruptions in the shapes and arrangements of most somatic muscles (Fig. 8F, cf. Fig. 8E). Only the VLMs, and particularly M12/VLM1 that expresses *tey* normally, are affected less severely (Fig. 8F and data not shown). Indeed, in many cases the genetic loss of *tey* function in M12p’s can largely be rescued by forced expression of *tey* via *tey-GAL4* (Fig. 8H, cf. Fig. 8G). Because the phenotypes obtained with the pan-mesodermal driver *twi-GAL4* did not allow a distinction between cell fate changes and migration defects as their cause, we utilized a later driver that is active in a small specific subset of muscle precursors, *org-1-GAL4* (Schaub et al., 2012). Ectopic *tey* using this driver also leads to mis-oriented and mis-shapen somatic muscles, which in this case sit in the background of unaffected muscles and still express the *org-1*-RFP and *slou*-LacZ markers used (Fig. 8J, cf. Fig. 8I). These data support our interpretation that ectopic expression of *tey* leads to migration and morphogenesis defects in somatic muscles rather than to cell fate transformations.

## Discussion

### Hallmarks of late-phase migrations of longitudinal visceral muscle precursors

The main focus of our study has been on the sparsely examined late phase of the migration of longitudinal visceral muscle precursors (LVMp’s), which occurs after these cells have completed their anterior migration and are spread out along the length of the trunk visceral mesoderm (TVM). This culminates in an intriguing developmental process that forms an orthogonal muscle pattern ensheathing the midgut. We show that during this process, these multinucleated muscle precursors perform very active migrations, with the net result being their dispersion towards the dorsal and ventral midlines, respectively, of the forming gut tube. It appears that these migrating LVMp’s attempt to attain the largest possible distance from their respective dorsal and ventral neighbors. Because the available space for migration continuously enlarges towards the dorsal and ventral sides until the closure of the midgut tube, the mutual distances of these migrating cells increase concomitantly, such that ultimately the longitudinal midgut muscles are distributed in an equidistant fashion around the circumference of the midgut.

This migratory behavior is reminiscent of the processes of cellular dispersion and cellular tiling, which has been described for several other migratory cell types (reviewed in Stramer and Mayor, 2017). A known example for cellular tiling is the even dispersal of *Drosophila* haemocytes along the ventral surface of the embryo (Davis et al., 2012). Similar to hemocyte tiling and other cellular dispersion processes, it is highly likely that the dispersion of LVMu’s over the expanding migration substrate involves mutual repulsion among the migrating cells. This notion is consistent with the observed increased distances of regularly spaced LVMu’s in mutants with reduced FGF activities, in which smaller numbers of surviving LVMp’s are present (Reim et al., 2012). Mutual repulsion is a major component during the process of Contact Inhibition of Locomotion (CIL), which has become a well-established phenomenon of cell migration (Stramer and Mayor, 2017). The distinct cellular steps and sub-cellular re-organizations taking place during CIL, which lead to this “phenomenon of a cell ceasing to continue moving in the same direction after contact with another cell” (Abercrombie, 1979), recently have been described in some detail (Stramer and Mayor, 2017; Roycroft and Mayor, 2018). Because the specific processes of CIL are known to be modified in different migrating cell types, which leads to distinct migration behaviors, it needs to be examined to which degree any of these are shared with the migrating LVMp’s.

We showed that the dorsoventral dispersion of the LVMp’s is accompanied by the presence of highly dynamic filopodial protrusions around the entire surface of these syncytia, which make transient contacts with filopodia or cell bodies of neighboring LVMp’s. For the migration of murine leukocytes, evidence has been presented suggesting that their lamellipodial and filopodial protrusions are largely functioning as ‘sensory organelles’ during migration (Leithner et al., 2016), and we propose that the filopodia of the migrating LVMp’s play analogous roles to mediate mutual repulsion among these cells. Of note, the prominent presence of filopodia between neighboring cells that do not show any directional bias and the lack of lamellipodia in LVMp’s are features that are also seen in migrating nascent myotubes in the developing *Drosophila* testes, which represent precursors of a different type of visceral muscle (Rothenbusch-Fender et al., 2017). A recent study showed that this latter process also involves CIL-related migration behavior. Detailed analyses indicated that the dynamics of the filopodial protrusions of these myotubes requires their formin-dependent formation and Rho1-linked retraction, as well as their Arp2/3 dependent branching and integrin-dependent cell-matrix adhesions, all of which appear to contribute to the normal migration of these cells (Bischoff et al., 2021). However, an important difference is that unlike LVMp’s, migrating testes myotubes maintain cohesiveness and migrate collectively towards the unoccupied space, which is more similar to the behavior of other collectively migrating cells. This difference could be reflected in relatively long-lived polarities from the cell-cell edges towards the free edges of the migrating testes myotubes versus highly transient repolarizations that may occur following the collisions of LVMu’s via their filopodial extensions. Therefore, these two migrating types of visceral muscle precursors must differ in some important aspects of their specific molecular interplays, even though parts of the molecular dynamics described by Bischoff et al. (2021) for the filopodia of testes myotubes are likely to be active in those of the LVMu’s as well.

A unique and intriguing feature of the migrating LVMp’s is that, during their dispersion over the midgut, they initiate mutual head-to-tail contacts with their closest anterior and posterior neighbors and align to form the longitudinal visceral muscles, each one of which extending over much of the length of the midgut. Hence, towards the end of their migration process, the proposed mutually repulsive activity initially present all around their surface must be suppressed at the anterior and posterior ends of these syncytia such that these ends can come into contact and establish stable adhesions. Although it is unclear how such a hypothesized locally-restricted switch from repulsive to adhesive properties within the syncytia might be accomplished, it must rely on pre-existing anisotropies within these cells. We believe that this anteroposterior anisotropy is already established in the context of the previous anterior migration of the LVMp’s as a loose collective, when they establish anterior-posteriorly elongated spindle shapes. The elongated shape of the LVMp’s that contain linear rows of nuclei is then maintained and even increased upon continued myoblast fusion and their dorsoventral migrations, and may rely on some of the same molecular events known to organize cytoskeletal components within somatic myotubes (Schulman et al., 2015). Even though some of these elongated migrating syncytia transiently can assume oblique or even transverse orientations (especially at midgut constrictions), during and after making contacts with their anterior and posterior neighbors they straighten along the anteroposterior axis and thus form the longitudinal midgut muscles (which appear to involve subsequent inter-syncytial fusion events, as indicated via clonal cytoplasmic markers (Klapper et al., 1998)). Although speculative, the strict dorsoventral orientation of the circular visceral muscle precursors, which serve as the migration substrate for the LVMp’s, could also influence the overall anteroposterior orientation and anisotropy of the migrating LVMp’s.

### Potential functions of Tey during visceral and somatic muscle precursor migration

In a previous study, our colleagues with some of us (A.I. & M.F.) reported on the role of Tey in the migration and synaptic targeting of motoneuronal axons (Inaki et al., 2010). Specifically, it was shown that the expression of Tey in the somatic muscle M12 serves to repress transcription of the Toll gene, which encodes a transmembrane receptor that therefore becomes restricted to the ventrally adjacent longitudinal muscles M13, M6, and M7 (aka, VLM2-4). Because Toll repels synapse formation of axons from the respective motoneurons, the repression of *Toll* by Tey in M12, together with the presence of this repulsive cue in the ventrally adjacent muscles that are crossed by the migrating axons, permits the specific neuromuscular targeting of M12. These and other data showed that the nuclear factor Tey can function as a transcriptional regulator.

Could Tey dependent *Toll* repression also be relevant for the functions of Tey during migration of visceral and somatic muscle precursors? In contrast to the effects of *tey* mutations, ectopic expression of *Toll* in M12 did not seem to perturb the localization and orientation of this muscle (Inaki et al., 2010), which would argue against a role of *Toll* repression by Tey in myotube attachment site targeting. However, as shown herein, the absence of *tey* function already affects the migration of M12 founder cells shortly after they are born, when the Mhc enhancer used for ectopically expressing Toll is not yet active. We think it is likely that mis-migration of this muscle founder in *tey* mutants contributes to, or is the underlying cause of the observed mis-positioning and aberrant attachment of M12, although we can not exclude a second requirement for *tey* in the specific targeting of the attachment sites by the growing myotube. Tey-dependent *Toll* repression still could possibly be involved in guiding M12 founder cell migration, but it seems more likely that Tey regulates genes encoding other cell surface or cytoplasmatic proteins guiding the migrating M12 founder cells. Likewise, our notion that the dispersion of the LVMp’s over the developing midgut is driven by mutual repulsion rather than by attraction to a specific target area (or lack of repulsion by the area, as during Tey-dependent axonal pathfinding) points to the involvement of different Tey downstream genes during this process. The observed phenotypes in *tey* mutants, where LVMp’s and their descendent longitudinal gut muscles fail to obey the normal distances from their lateral neighbors and often touch them, suggests that Tey could regulate the expression of some of the genes involved in CIL, as discussed above. Additionally, the observation that the LVMp’s are less well oriented in anteroposterior directions and generally fail to align with their anterior and posterior neighbors upon finishing their migration makes it conceivable that Tey regulates some of the yet unknown activities involved in cell anisotropy and mutual adhesion of myotube tips as well. More broadly, we propose that the role of Tey in regulating the migration of these muscle precursors is somewhat analogous to the role of Slow border cells (Slbo), a cell type-specific transcription factor that is thought to regulate various migration-related target genes necessary for the migration of border cells during *Drosophila* ovary development (Montell et al., 2012).

In addition to its role in LVMp migration and alignment, *tey* is required for proper morphogenesis and differentiation of longitudinal visceral muscles (LVMu’s), as is most obvious from the highly abnormal shapes of these muscles and the aberrant lateral myofibril alignments within them in larval and adult midguts from *tey* mutants. For somatic muscles in *Drosophila* it was shown that lateral myofibril alignment requires mechanical tension caused by twitching during late stages of their differentiation (Weitkunat et al., 2017). Thus, the myofibril mis-alignments in LVMu’s from *tey* mutants in part may be a result of reduced mechanical tension because of the failed head-to-tail alignments of the migrating LVMp’s. Alternatively (or perhaps additionally), this phenotype could reflect direct functions of *tey* in LVMu differentiation that are independent of its earlier requirements in LVMp’s and involve yet other sets of downstream genes.

### How does Tey function molecularly?

As Tey lacks any DNA binding domain and is homologous to the vertebrate E3 ubiquitin ligase RNF220 it must exert its gene regulatory functions indirectly. Like Tey, murine RNF220 is expressed in neuronal tissues of the CNS, but unlike Tey it is not known to be expressed in developing muscle tissues. In various neuronal contexts, murine RNF220 has been documented to mono- or poly-ubiquitinate several transcription factors, chromatin regulators, and nuclear effectors of the Wnt and Sonic Hedgehog pathways (reviewed in Ma and Mao, 2022). For some of these factors, RNF220-dependent ubiquitination has activating or stabilizing effects whereas for others it leads to their inactivation or degradation (Ma and Mao, 2022). Many of these functions require the RNF220 binding partner ZC4H2 (a C4H2-type zinc finger protein) and mutations in the genes for the two proteins cause similar neuronal phenotypes (Ma and Mao, 2022). Interestingly, two independent large-scale protein interaction screens found the *Drosophila* ortholog of ZC4H2, CG13001, to bind to the paralog of Tey, CG4813 (Guruharsha et al., 2011; Hu et al., 2017), which may indicate that some of the molecular functions of murine and *Drosophila* RNF220 proteins are shared. However, because the functions of Tey described herein and in Inaki et al. (2010) concern developing muscle tissues, and because the RING finger domain of Tey has strongly diverged, its relevant molecular targets in these tissues are likely to be different. Their identities and functions, as well as the questions of whether Tey requires CG13001 binding and functions as an E3 ubiquitin ligase despite its diverged C-terminus, need to be addressed in future studies.

## Acknowledgements

We gratefully acknowledge the important contributions of the following colleagues: Sharon Israeli for sequencing *tey* cDNAs prior to the completion of the genome project, Johannes März for the stainings shown in Fig. 5E and Fig. 7A-D, and Christoph Schaub for teaching MF various CRISPR/Cas9 and cloning techniques. We thank Renate Renkawitz-Pohl, Alexandra Schambony, and the Developmental Studies Hybridoma Bank (DHSB, Univ. of Iowa) for providing antibodies and the Bloomington Stock Collection (Univ. of Indiana), as well as the colleagues listed in Materials & Methods, for providing fly stocks. We also thank Michael Schoppmeier and Wiebke Herzog for providing postretirement lab space to MF.

## Movies

**Movie 1.** Time lapse movie of LVMp and LVMu migration in heterozygous *tey^5053A^* control embryo from mid stage 14 to stage 17. LVMp’s are marked with tey > *lifeact-GFP* (green) and CiVMp’s with *bap3-RFP* (magenta). Open arrow heads: Examples of filopodial extensions making dynamic contacts between neighboring LVMp’s and LVMu’s. Closed arrow heads: Differentiating LVMu’s at st. 17. M12: Somatic muscle M12.

**Movie 2.** Time lapse movie of LVMp migration in homozygous *tey^5053A^* mutant embryo from early stage 14 to stage 16 (marked and labeled as in Movie 1).

## Supplemental Figures

**Fig. S1. Lack of Tey protein expression in *tey^5053A^* homozygous embryos and congruence of *tey*-GAL4 activity with Tey expression in heterozygotes.** (A, A’) Homozygous st. 14 *tey^5053A^ UAS-lacZ* embryo showing reporter expression in longitudinal visceral and somatic M12 muscle precursors (A) but no Tey protein in these or any other cells. (B, B’) Embryo at early st. 14 *tey^5053A^ UAS-lacZ* in trans to *TM3 eve-LacZ* balancer showing spatial congruence of *tey*-driven reporter gene activity and nuclear Tey protein in longitudinal visceral and somatic M12 muscle precursors (as well as striped balancer-derived βGal expression).

**Fig. S2. Sequence alignments of wildtype and mutant versions of Tey from *Drosophila melanogaster* with its orthologous proteins from *Aedes egypti, Tenebrio molitor, mus musculus*, and its *D. mel*. paralog CG4813.** The highly conserved RNF220 domains and RING domains are boxed in magenta and green, respectively. The bars on top of the sequences mark stretches with predicted α-helical (light blue), looped (dark blue) and β-sheet (green) conformations in fly Tey and mouse RNF220, respectively, as derived from the AlphaFold Protein Structure Database. (A stretch indicated by the thin line between brackets is absent in the mouse isoform used in AlphaFold). The C-termini of the mutant Tey^Δ.M2-1^ and Tey^Δ.M1-11^ proteins are depicted in red on top of the Tey (*D. mel*.) sequence. The amino acids preceding the vertical bars represent the last residues of native Tey present in the respective mutant versions, which in Tey^Δ.M2-1^ is followed by a stop codon and in Tey^Δ.M1-11^ by a short out-of-frame peptide sequence. Likewise for *Tey^ΔRNF.sfGFP^* the transition between the N-terminal Tey sequences and a linker plus sfGFP is indicated.

**Fig. S3. Longitudinal visceral muscle phenotypes in embryos with various *tey* allelic combinations.** Left hand column shows controls and right hand column *tey* mutants. (A – D) Stage 14 and stage 16 embryos, respectively, carrying *HLH54F-GFP* together with *tey^Δ.M1-11^* in trans to *TM6 Dfd-EYFP* (left) or *Df(3L)ED228* (right). Anti-GFP is shown in green and anti-Fas3 in magenta. (E – H) Stage 15 and stage 16 embryos, respectively, carrying *HLH54F-cyto-RFP* together with *tey^ΔRNF.sfGFP^* in trans to *TM6 Dfd-EYFP* (left) or *Df(3L)ED228* (right). Anti-GFP is shown in green and anti-RFP in magenta. (I, J) Stage 16 embryos carrying *UAS-lifeact-GFP* in trans to *bap3-RFP* on the second chromosome together with *tey^5053A^* in trans to *TM6 Dfd-EYFP* (left) or *Df(3L)ED228* (right). Anti-GFP is shown in green and anti-RFP in magenta. (K, L) Stage 15 embryos carrying *UAS-apoliner9* on the second chromosome together with *tey^5053A^ UAS-lacZ* in trans to *TM6 Dfd-EYFP* (left) or *Df(3L)ED228* (right). Anti-RFP is shown in red, anti-βGal in green, and Hoechst-stained DNA in blue. As in the control (K), no nuclear RFP is detectable in the mutant (L), which argues against increased apoptosis in longitudinal visceral muscle precursors lacking *tey* function (a and p denote anterior and posterior, respectively). (M – N’) Stage 15 embryos with *tey^ΔRNF.sfGFP^* in trans to *TM6 Dfd-EYFP* (left) or *Df(3L)ED228* (right) were stained with anti-GFP for visualizing TeyΔRNF::sfGFP and with anti-Tey (magenta; ca. 50 % nuclear in the heterozygous control and largely cytoplasmatic in the mutant). For better clarity, the phenotypes in the longitudinal visceral muscle precursors were separated from those of the somatic M12 muscle precursors in the same embryos by showing the Z-planes of the former in (M, N) and the Z-planes of the latter in (M’, N’). Scale bars provided in control panels also apply to the other panels with controls or mutants from the respective series.

**Fig. S4. *tey* phenotypes in midguts of 3^rd^ instar larvae (high magnifications) and adult flies**. (A, B) 3^rd^ instar larval midguts with *tey^5053A^ UAS-lacZ* in trans to *TM6 Dfd-EYFP* (A) or *Df(3L)A23* (B) and stained with anti-βGal for visualizing the longitudinal visceral muscles and for F-actin (Alexa Fluor™ *555* Phalloidin) in all gut muscles. Examples of nuclei within individual longitudinal visceral muscle fibers (visible due to partially nuclear GFP) are marked by arrow heads, which in the mutant allows the assignment of individual, separated actomyosin fibrils to a distinct muscle cell. (C, D) 3^rd^ instar larval midguts with *UAS-lifeact/bap3-RFP* together with *tey^5053A^* in trans to *TM6 Dfd-EYFP* (C) or *Df(3L)ED228* (D) and stained with anti-GFP for visualizing the sarcomeres of longitudinal visceral muscles and for F-actin (Alexa Fluor™ *555* Phalloidin) in all gut muscles (anti-RFP is omitted as it was negative in larvae). (E, F) Midguts from adult wildtype (E) and *tey^5053A^/Df(3L)ED228* escaper fly stained for F-actin (Alexa Fluor™ *555* Phalloidin). The mutant gut (same as in Fig. 5K) has a chambered and thickened appearance, which appears to result from its excessive contraction by the aberrantly-attached longitudinal gut muscles along the a/p axis.

**Fig. S5. Guide RNA sequences and primers used for generating CRISPR/Cas9 induced *tey* mutants.**

## Supplemental Movies

**Movie S1:** Time lapse movie of LVMu migration in heterozygous *tey^5053A^* control embryo from mid stage 16 to stage 17 (focusing on late events; marked and labeled as in Movie 1).

**Movie S2:** Time lapse movie of LVMp and LVMu migration in homozygous *tey^5053A^* mutant embryo from mid stage 14 to stage 17 (mainly focusing on late events; marked and labeled as in Movie 1).

